# Modeling synaptic integration of bursty and beta oscillatory inputs in ventromedial motor thalamic neurons in normal and parkinsonian states

**DOI:** 10.1101/2023.04.14.536959

**Authors:** Francesco Cavarretta, Dieter Jaeger

## Abstract

The Ventromedial Motor Thalamus (VM) is implicated in multiple motor functions and occupies a central position in the cortico-basal ganglia-thalamocortical loop. It integrates glutamatergic inputs from motor cortex (MC) and motor-related subcortical areas, and it is a major recipient of inhibition from basal ganglia. Previous experiments in vitro showed that dopamine depletion enhances the excitability of thalamocortical cells (TC) in VM due to reduced M-type potassium currents. To understand how these excitability changes impact synaptic integration in vivo, we constructed biophysically detailed VM TC models fit to normal and dopamine-depleted conditions, using the NEURON simulator. These models allowed us to assess the influence of excitability changes with dopamine depletion on the integration of synaptic inputs expected in vivo. We found that VM TCs in the dopamine-depleted state showed increased firing rates with the same synaptic inputs. Synchronous bursting in inhibitory input from the substantia nigra pars reticulata (SNR), as observed in parkinsonian conditions, evoked a post-inhibitory firing rate increase with a longer duration in dopamine-depleted than control conditions, due to different M-type potassium channel densities. With beta oscillations in the inhibitory inputs from SNR and the excitatory inputs from drivers and modulators, we observed spike-phase locking in the activity of the models in normal and dopamine-depleted states, which relayed and amplified the oscillations of the inputs, suggesting that the increased beta oscillations observed in VM of parkinsonian animals are predominantly a consequence of changes in the presynaptic activity rather than changes in intrinsic properties.

**Significance Statement:** The Ventromedial Motor Thalamus is implicated in multiple motor functions. Experiments in vitro showed this area undergoes homeostatic changes following dopamine depletion (parkinsonian state). Here we studied the impact of these changes in vivo, using biophysically detailed modeling. We found that dopamine depletion increased firing rate in the ventromedial thalamocortical neurons and changed their responses to synchronous inhibitory inputs from substantia nigra reticulata. All thalamocortical neuron models relayed and amplified beta oscillations from substantia nigra reticulata and cortical/subcortical inputs, suggesting that increased beta oscillations observed in parkinsonian animals predominantly reflect changes in presynaptic activity.

## Introduction

Motor thalamus is a critical structure for multiple aspects of motor control (Bosch-Bouju *et al*., 2013). In rodents, it comprises the ventro-medial (VM), the ventral-anterior (VA), and the ventro-lateral nuclei (Kuramoto *et al*., 2011). In particular, VM and VA are the major recipients of inhibition from substantia nigra pars reticulata (SNR) (Chevalier and Deniau, 1982; Kuramoto *et al*., 2011; Rovó *et al*., 2012), the output area of the basal ganglia. Excitatory inputs from primary and pre-motor cortices (MC) play a key role in VM processing, supporting persistent motor preparatory activity in a closed loop (Guo et al., 2017; Guo et al., 2018). In vivo experiments implicated VM in movement preparation and vigor control (Takahashi *et al*., 2021), while SNR mediates strong and temporally precise inhibition that controls movement direction and initiation in behaving animals (Morrissette *et al*., 2019; Catanese and Jaeger, 2021).

Parkinson’s disease (PD) is a neurodegenerative disorder that is primarily due to the death of dopamine neurons in the Substantia Nigra pars compacta (SNC). In rodents, dopamine depletion increases synchrony and bursting in the SNR (Lobb and Jaeger, 2015; Brazhnik *et al*., 2016; Willard *et al*., 2019). This maladaptive activity is conveyed to VM, and this pathway may be associated with deficits in movement selection and initiation (Kravitz *et al*., 2010; Morrissette *et al*., 2019; Takahashi *et al*., 2021; Tekriwal *et al*., 2021). Moreover, local field potential recordings showed increased beta oscillations in SNR, MC, and VM of parkinsonian rats (Brazhnik et al., 2012; Brazhnik et al., 2016; Nakamura et al., 2021), a typical hallmark of parkinsonian pathophysiology. In our previous publication, we studied the effects of dopamine depletion on the VM of mice. Our results showed enhanced excitability of thalamocortical neurons (TC), with increased rebound bursting upon hyperpolarization (Bichler *et al*., 2021). While experimental evidence suggests that the nigral synapses are well positioned to evoke rebound bursting in VM TCs (Bodor *et al*., 2008; Edgerton and Jaeger, 2014b; Bichler *et al*., 2021), this hypothesis has not been tested in vivo. It remains unknown whether these effects of dopamine depletion on intrinsic VM properties contribute to the generation of beta oscillations in PD (Brazhnik *et al*., 2016), or exacerbate rebound bursting in vivo.

Because the contribution of intrinsic properties to firing patterns in vivo is hard to measure directly, we employed biophysically detailed modeling, using the NEURON simulator (Hines and Carnevale, 1997). Specifically, we fitted TC models, replicating the different firing properties observed in normal and parkinsonian states (Bichler *et al*., 2021). We implemented the properties of afferent inputs to VM, replicating synaptic conductances and subcellular distributions observed experimentally (Bodor *et al*., 2008; Edgerton and Jaeger, 2014b; Gornati *et al*., 2018a), where the synapses were activated by artificial spike trains replicating the activity during motor performance (Inagaki *et al*., 2022). This approach allowed us to control the firing patterns of the inputs, to reproduce either normal firing features or distinct firing features of the parkinsonian state. We then added varying amounts of synchronous bursting or beta oscillations to these inputs. We found that TCs in dopamine-depleted state responded at higher firing rate than TCs in normal conditions to all in vivo like input patterns. Synchronous nigral bursting was unable to evoke rebound bursting, as the synaptic excitation prevented sufficient hyperpolarization to de-inactivate T-type Ca^2+^ channels. However, synchronous nigral bursts still evoked a significant post-inhibitoryfiring rate increase in TCs, due to slow recovery of potassium currents upon repolarization. This increase lasted longer in models fit to dopamine-depleted conditions. Adding beta oscillations in SNR inputs resulted in significant spike-phase locking in TC firing in both normal and parkinsonian states. This phase locking became stronger when excitatory inputs contained beta oscillations at a 180-degree phase shift. These results suggest that the homeostatic changes induced by dopamine depletion do not affect the beta oscillations in the VM, while VM TCs in both states are able to amplify such oscillations.

## Materials and Methods

### Simulation

The morphology of a reconstructed thalamocortical neuron was divided in 40 μm-long compartments and used to construct multicompartmental biophysically detailed models with 11 Hodgkin-Huxley-style (HH) active conductances at varying densities (for details, see *“Neuron morphology”* and *“Intrinsic membrane properties”*).

Simulations were implemented in a fully integrated NEURON and Python environment (Hines and Carnevale, 1997). To fit the neuron models to physiological targets, we employed multi-objective optimization, based on genetic algorithms, using the BluePyOpt package (Van Geit *et al*., 2016). Here each solution corresponds to a neuron model, encoded as an array of parameters associated with the intrinsic properties. The optimizer calculated model fitness by comparing the traces generated by each model against a set of features extracted from experimental recordings, simulating a battery of experiments with adaptive time steps, using the CVode solver for partial differential equations (for more details, see *“Fitting neuron models”)*. The fitting process was executed on the supercomputer Expanse, managed by the San Diego Super Computer Center (SDSC), through the Neuroscience Gateway (NSG; http://www.nsgportal.org. Sivagnanam *et al*, 2013). Following model construction, simulations to evaluate model performance were executed using the NEURON version known as “CoreNEURON” (Kumbhar *et al*., 2019), and parallelized with the MPI4Py package. These simulations were executed on the supercomputer Expanse.

All simulation parameters such as temperature, reversal potentials of ion species, effects of ion channel blockers, and holding membrane voltage reflected the specifics of the experimental stimulation protocols and slice conditions (Table 1), while the number of active synapses reproduced experimentally observed values (Table 1). For each experiment, we calculated the values of reversal potential for sodium and potassium using the Nerst equation, accounting for the relative concentrations of ions in the artificial cerebrospinal fluid (aCSF) and pipette solutions. In some experiments, tetradoxine (TTX), a sodium channel blocker, was applied, and/or the pipette solution contained cesium, a potassium channel blocker. We accounted for their effects by turning off sodium and potassium HH-conductances in the simulations, respectively.

**Table 1.** Simulated experimental conditions. In this work, the simulations were replications of real experiments, accounting for temperature, holding membrane voltage (V_hold_), and reversal potential of ion species (E_rev_), such as sodium (Na), potassium (K), and chloride (Cl). The holding membrane voltages were corrected for the junction potential.

| Reference | Temp. (°C) | V<sub>hold</sub> (mV) | Experimental conditions |
| --- | --- | --- | --- |
| Bichler et al, 2021 | 27 | -69 | E<sub>Na</sub>: 69 mV<br>E<sub>K</sub>: -105 mV |
| Laudes et al, 2012 | 24 | -79.3 | TTX application and pipette filled with cesium blocking sodium and potassium channels, respectively |
| Gornati et al, 2018 | 34 | -68.4 | TTX application blocking sodium channels<br>E<sub>K</sub>: -107.1 mV |
| Edgerton and Jaeger, 2014 | 32 | -64 | E<sub>Na</sub>: 60.1 mV<br>E<sub>K</sub>: -105.8 mV |

### Code Accessibility

The source code is publicly available on ModelDB and GitHub (https://github.com/FrancescoCavarretta/BGMT).

### Modeling process overview

We designed a data-driven modeling pipeline (Fig. 1) to build realistic models of thalamocortical neurons (TC) from ventromedial motor thalamus (VM) of mice in normal and parkinsonian states (green boxes) along with afferent inputs (blue boxes). We defined a biophysically detailed model for TCs, comprising a full three-dimensional morphology from the VM of mouse (see *“Neuron morphology”*) and 11 HH models of ion channels (see *“Intrinsic membrane properties”*), with subcellular distributions based on experimental data of thalamic and pyramidal cortical neurons (see *“Ion channel distributions”*). The model comprised multiple free parameters, with values determined by multi-objective optimization (see *“Template data taken from in vitro experiments”* and *“Fitting neuron models”*). We then modeled the main synaptic inputs to VM, reproducing the subcellular distributions, numbers of synaptic contacts, and unitary conductances observed experimentally (see *“Modeling synaptic inputs”*), with realistic levels of presynaptic activity (see *“Pre-synaptic activity: artificial spike train generation”*).

**Figure 1.**
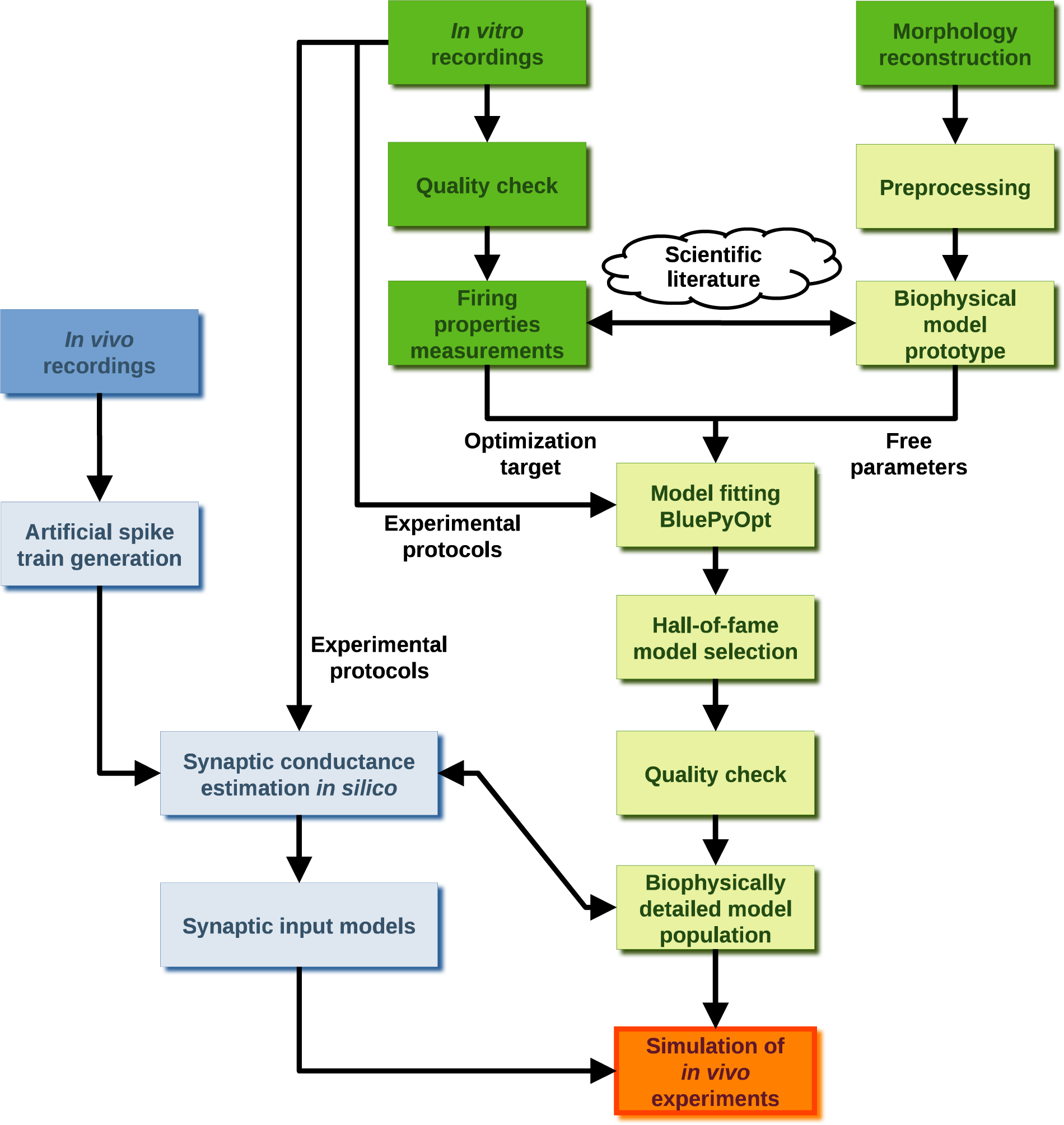
Simulation of in vivo firing activity of ventromedial motor thalamus in normal and parkinsonian states. The data-driven modeling pipeline for biophysically detailed simulations of in vivo conditions for thalamocortical neurons of ventromedial motor thalamus in normal and parkinsonian states. The pipeline comprises two branches, single cell (green) and synaptic afferent (blue) modeling, which converge on a single aim, the simulation of in vivo activity of mouse ventromedial motor thalamus in normal and parkinsonian states (orange). Both branches start from the analysis of anatomical and electrophysiological data (dark green and dark blue), integrated with the information reported in literature (cloud).

### Neuron morphology

We based our model on a full three-dimensional reconstruction of a thalamocortical neuron from the ventromedial thalamic nucleus (VM) of mouse (id. AA0719;https://www.janelia.org/project-team/mouselight). For modeling purposes, we retained only the proximal 70 μm-long portion of the axon (i.e., axonal initial segment), approximated by two cylindrical compartments (diameter: 1.5 μm; length: 35 μm), as we did not further follow action potential propagation. The original reconstruction lacked estimates for dendritic diameters while the soma was represented by the cross-sectional perimeter only. We replaced the soma with a cylindrical section (diameter: 36.9 μm; length: 24 μm), yielding a cross-section surface of 293 μm^2^, consistent with the experimental estimates of VM thalamocortical cells (TC) of rats (cf 288±13 μm^2^ and 298±2 μm^2^; ±SEM) (Sawyer *et al*., 1989). To model the dendritic diameters, we defined a recursive formula based on Rall’s 2/3 Power Law:
- If the dendrite has children

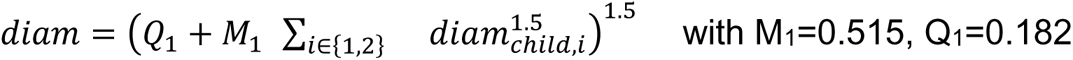

- otherwise

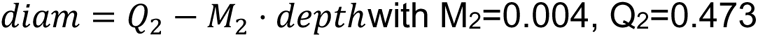

where *diam* indicates the dendritic diameter, *depth* indicates the number of branch points from soma, and *diam_child,i_* is the diameter of the i^th^ child. Additionally, for primary dendrites only, the diameter tapers with the distance from soma (*d*):

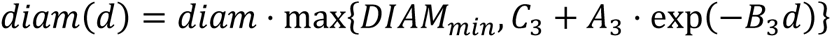

with A_3_=0.861, B_3_=0.045, C_3_=0.858, DIAM_min_=1 with *diam* defined as above. Figure 3 (B) shows the resulting dendritic diameters. The values of the parameters, i.e., *M_*_, Q_*_, A_3_, B_3_, C_3_*, and *DIAM_min_*, were estimated by calculating the least square regression curves from 5 partial reconstructions of VM thalamocortical cells of mouse obtained in our laboratory.

### Intrinsic membrane properties

NEURON models require the specification of passive properties, such as specific membrane resistivity (r_m_), axial resistivity (r_i_; ie, intracellular or cytoplasmic resistivity), specific membrane capacitance (c_m_), and resting potential (V*_Rest_*). In our models, their values were uniform across all sections. In particular, c_m_ was a constant using the standard canonical value (1 μF/cm^2^) (Gentet *et al*., 2000) while the others were treated as free parameters estimated by our genetic algorithm (see *“Fitting neuron models”*). The active membrane properties of our models consisted of 11 HH-conductances, with states dependent on the membrane potential and/or intracellular Ca^2+^ concentration. They were transient (Na_T_) and persistent (Na_P_) sodium currents; delayed rectifier (K_DR_), A-type (K_A_), delaying (K_D_), and M-type (K_M_) potassium currents; H-type nonspecific cation current (I_H_); T-(Ca_T_) and L-(Ca_L_) type Ca^2+^ currents; small conductance (SK) and big conductance (BK) Ca^2+^-dependent potassium currents. Most HH-conductances were specified as used in a previous model of ventrobasal TC (Na_T_, K_DR_, K_A_, I_H_, Ca_T_, Ca_L_, SK) (Iavarone *et al*., 2019), while K_M_ was specified as used in our previous model of VM TC (Bichler *et al*., 2021). We derived the Na_P_ dynamics from the Na_T_ model (Iavarone *et al*., 2019), decreasing the half-values of activation and inactivation curves by 14 mV, consistent with experimental estimates from pyramidal cells (Hu *et al*., 2009). We based the K_D_ model on the dynamics of the shaker-related potassium channels observed in TC neurons of rats (Lioudyno *et al*., 2013), where the effects of these channels delayed the AP onset (Lioudyno *et al*., 2013), similarly to the effects of K_D_ currents observed in pyramidal neurons (Storm, 1988). We added the BK to induce firing rate adaptation in TCs (Ehling *et al*, 2013), observed also in VM (Fig. 2) (Bichler *et al*., 2021). The BK dynamics were voltage and Ca^2+^-dependent (Rothberg and Magleby, 2000), and accounted for the dependence on temperature (Q_10_=2.5) (Yang and Zheng, 2014).

**Figure 2.**
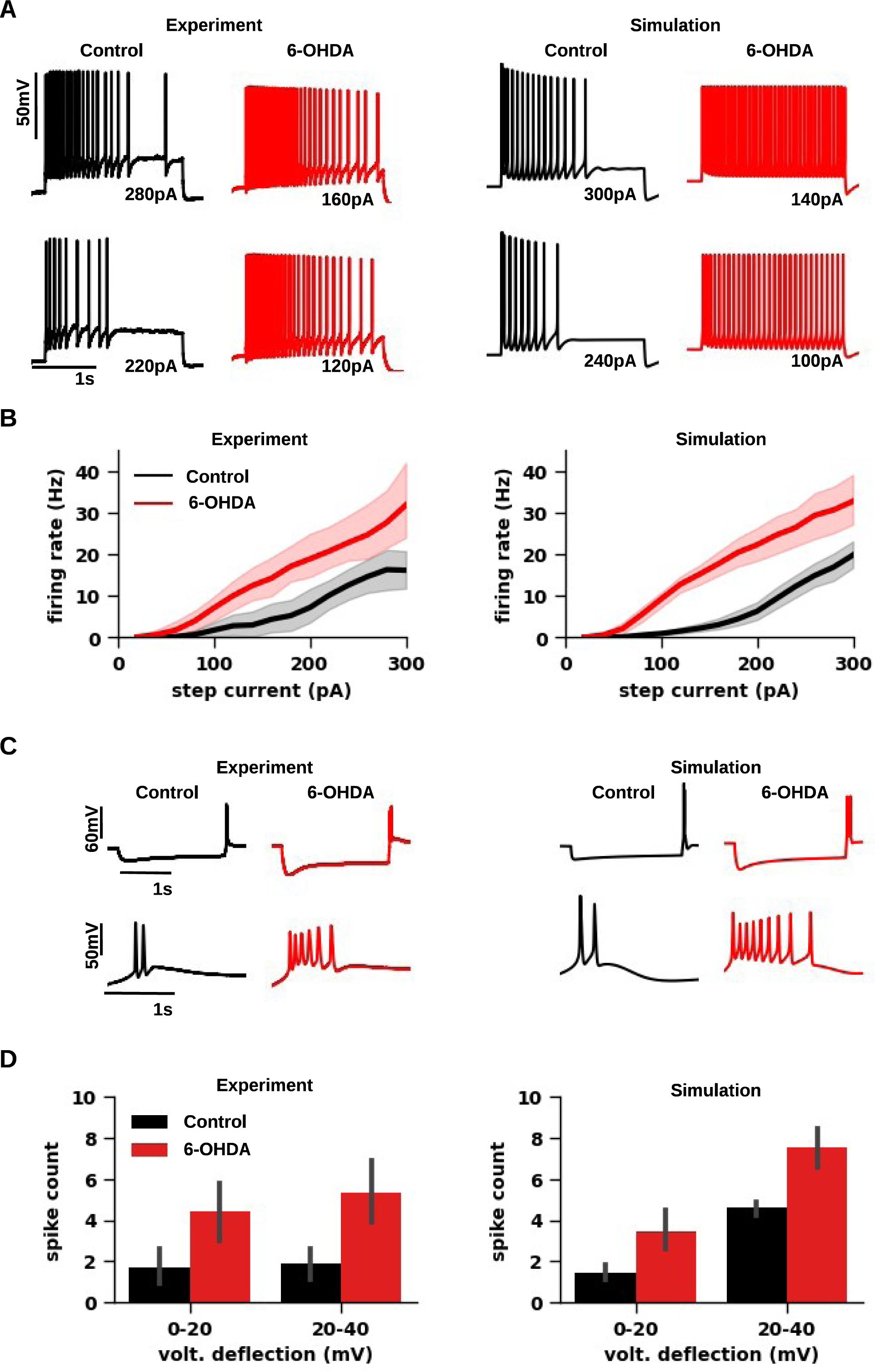
Experimental and simulated responses of thalamocortical neurons from ventromedial motor thalamus in normal and parkinsonian states. ***A***, Representative neuron responses to increasing depolarizing current injections. The baseline membrane potential was depolarized to −69 mV by a bias current (control: 93 pA and 6-OHDA: 67 pA). The bias current inactivated T-type Ca^2+^ channels and thus enabled tonic firing generation. ***Left***, Whole-cell recordings from ventromedial thalamus of control mouse (black) and 20 weeks after 6-OHDA injection (red). ***Right***, Simulated neuron responses of thalamocortical neurons from ventromedial thalamus of mouse in normal (black) and parkinsonian (red) models. ***B***, The f-I curves for thalamocortical neurons of ventromedial motor thalamus in normal (black) and parkinsonian (red) states. Dopamine depletion increased the firing rate and shifted f-I curves to the left (Wilcoxon, p < 0.001) in both experiments (control: n=9 and 6-OHDA: n=7 cells; left) and simulations (control: n=17 and 6-OHDA: n=12 models; right). The step current (20-300 pA with 20 pA increments) was delivered on top of the bias current (same as in ***A***). ***C,*** Representative neuron responses to hyperpolarizing current injection (−200 pA) on top of a bias current (same as in ***A***), evoking rebound bursting at the offset of the step current. ***Left***, Whole-cell recordings from ventromedial thalamus of control mouse (black) and after 6-OHDA treatment (red). ***Right,*** Simulation of the same stimulation protocol and experimental conditions with normal (black) and parkinsonian (red) neuron models. ***D,*** Bar graph pairs representing the average rebound spike count (±SEM) sorted by the amplitude of voltage deflection reached during hyperpolarizing current steps (−50 pA to –200 pA with 50 pA increments) in controls (black) and in parkinsonian (red) conditions. Rebound spike count was significantly higher for parkinsonian neurons than control in both experiments (Mann-Whitney, 20-40 mV: p=0.015; 40-60 mV: p=0.01; control: n=9 and 6-OHDA: n=7 cells; left) and simulations (Mann-Whitney, 20-40 mV: p=0.001; 40-60 mV: p=0.003; control: n=17 and 6-OHDA: n=12 models; right).

**Figure 3.**
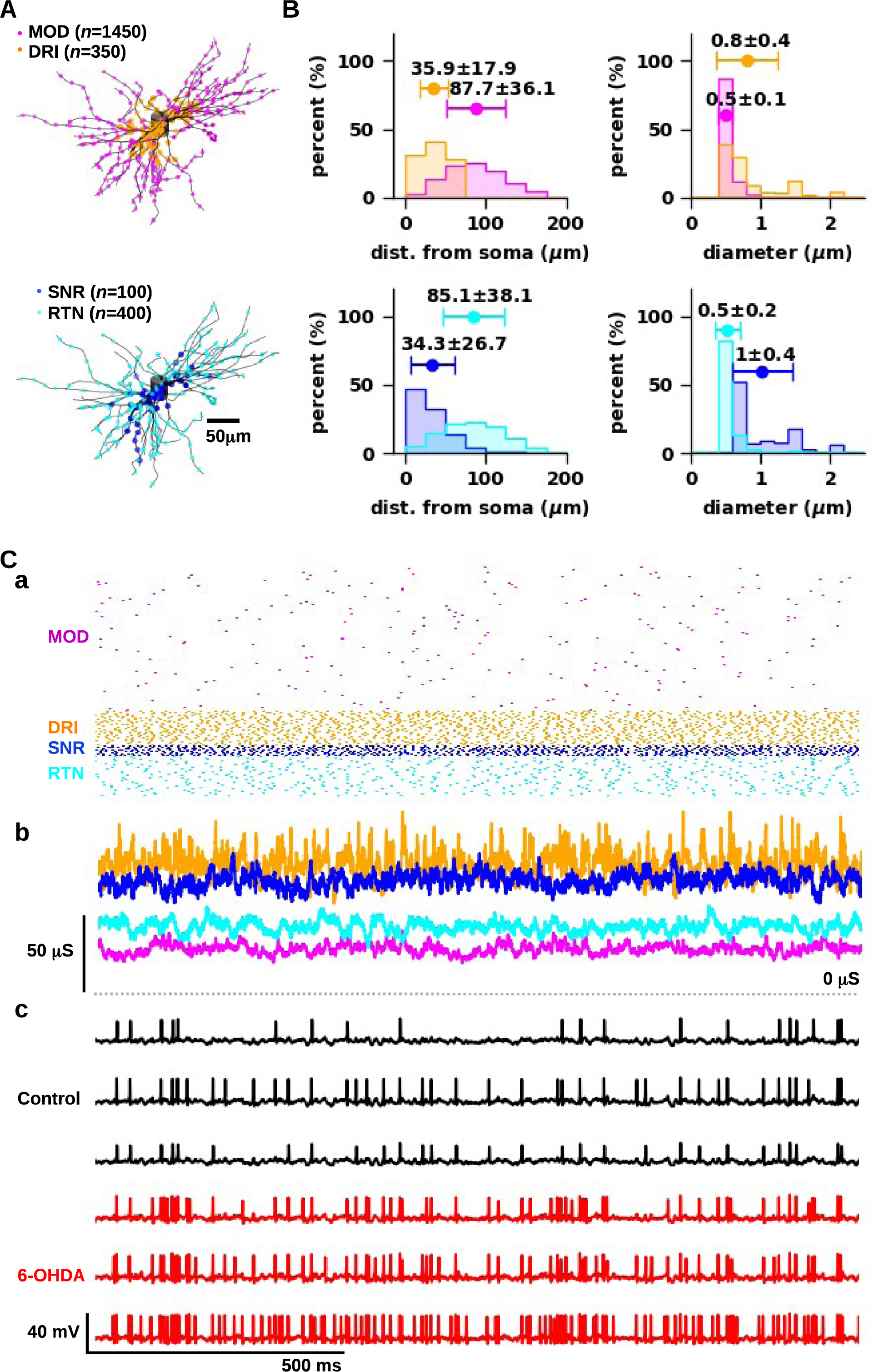
Simulation of normal and parkinsonian activity in vivo. ***A,*** Postsynaptic locations of synapses. ***Top,*** locations of excitatory modulators (MOD; magenta) and drivers (DRI; orange). ***Bottom,*** locations of inhibitory synapses from substantia nigra reticulata (SNR; blue) and reticular thalamic nuclei (RTN; cyan). ***B,*** Statistical distributions of distance from soma (left) and diameters (right) of the postsynaptic dendritic segment for excitatory (top) and inhibitory (bottom) inputs (as in A). ***C,*** Simulation of in vivo conditions. ***a,*** Representative artificial spike trains representing the spontaneous presynaptic activity with physiological values of firing rate (MOD: 1.1±0.1 Hz; DRI: 31.5±0.5 Hz; SNR: 52.4±0.7 Hz; RTN: 10.5±0.3 Hz; ±SD) and coefficient of variation of inter-spike intervals (0.43±0.06 for all). ***b,*** Total synaptic conductance for each set of synaptic inputs, as shown in *a*. ***c,*** Representative responses evoked by synaptic inputs to models of thalamocortical neurons in normal (black) and parkinsonian (red) states. The two states underlie the generation of different firing activity patterns.

Consistent with previous experimental studies (Womack *et al*., 2004), intracellular Ca^2+^ was structured in three separate microdomains, associated with distinct decay time constants. In particular, the Ca^2+^ flowing through Ca_L_ channels interacts with two microdomains that activate SK and BK channels separately, while Ca^2+^ flowing through Ca_T_ channels interacts with a third microdomain that does not activate other ion channels. In our models, the Ca^2+^ concentration of each microdomain determined the reversal potential for the related Ca^2+^ channel, calculated with the Goldman–Hodgkin– Katz flux equation.

### Ion channel distributions

Based on experimental data of thalamocortical neurons (Budde *et al*., 1998; Williams and Stuart, 2000), we modeled the subcellular distributions of ion channels as follows:

- The conductance density of voltage-dependent potassium channels (i.e., K_DR_, K_A_, K_M_, K_D_) is uniform throughout the somatodendritic shaft.
- The conductance density of sodium channels (i.e., Na_T_, Na_P_) decreases at each branch point.
- For T-type Ca^2+^ channels (i.e., Ca_T_), the conductance density is double as that of the soma for primary dendrites, and half of the soma for all the other dendrites.
- The conductance density of L-type Ca^2+^ channels (i.e., Ca_L_) decreases to half of the soma within the proximal 10μm-long portion of primary dendrites, and to a third in all other dendritic locations.

As the subcellular distribution of small- and big-conductance Ca^2+^-dependent potassium channels (i.e., Ca_L_, SK, BK) is unknown in TCs, and given the lack of data for the axon initial segment (AIS) of TCs, we modeled their distributions on data from pyramidal cells (Bowden *et al*., 2001; Hu *et al*., 2009; Battefeld *et al*., 2014):

- As Ca_L_ and the Ca^2+^-dependent conductance densities (i.e., SK) are co-localized in pyramidal cells, we used the same subcellular distributions for SK and BK, while these channels were absent in the axon.
- Na_T_ and Na_P_ conductance densities can be up to 19-fold higher than in the soma in the proximal and distal halves of the AIS, respectively.
- K_M_ conductance density in the distal half of the AIS can be up to 50 times higher than in the soma.

In the absence of specific information, we made the following assumptions:

- H-type conductance density is uniform along the somatodendritic arbor but absent in the AIS.
- K_DR_ conductance density can be up to 5-fold higher in the proximal half of the AIS than in the soma to compensate for the effects of the high Na_T_ conductance density, thus enabling full action potential repolarization.

### Template data taken from in vitro experiments

We selected a subset of whole-cell recordings from our previous study *in vitro* (*Bichler et al., 2021*), performed on adult mice in normal conditions and after unilateral 6-OHDA injections placed in the median forebrain bundle (normal: n=9; 6-OHDA: n=7). These recordings were obtained with different experimental protocols:

- Protocol 1: 2s-long depolarizing current steps (20-300 pA with 20 pA increments) on top of a bias current that held the membrane potential at −69 mV (normal: 93 pA; 6-OHDA: 67 pA), to estimate f-I curves.
- Protocol 2: 2s-long hyperpolarizing current steps (from –200 pA to –50 pA with 50 pA increments) on top of a bias current (same as in Protocol 1), to estimate the rebound spike count versus current relationship.
- Protocol 3: 2s-long hyperpolarizing steps (−200 pA to –50 pA with 50 pA increments; without bias current), to measure sag amplitude and evoke rebound bursting versus current relationships.
- Protocol 4: single pulse of –1 nA and 0.5 ms of duration, to measure membrane time constant.
- Protocol 5: 100 ms-long current step of –10 pA, to estimate input resistance.

To measure firing properties from simulation and experimental traces, we used a custom version of the eFEL package (https://github.com/BlueBrain/eFEL). In particular, the targets for model performance were constituted by membrane and firing properties extracted from our experimental recordings obtained with protocols (see Table 2 for a complete list).

**Table 2.** Summary of experimental protocols and targets for neuron model evaluation during the fitting process. Labels in square brackets indicate parameter name in source code.

|  |  |  |
| --- | --- | --- |
| **Protocol 1** | **Control Condition**<br>Steps: 200 pA, 240 pA, 280 pA<br>Bias: 93 pA | **Parkinsonian Condition**<br>Steps: 120 pA, 180 pA, 240 pA<br>Bias 67 pA |
| Avg. action potential (AP) amplitudes [AP_amplitude] |  |  |
| Avg. slow after-hyperpolarization (AHP) depth [AHP_depth] |  |  |
| AP_width [Avg. AP width] |  |  |
| AP_count |  |  |
| First-spike latency |  |  |
| Instantaneous firing rate calculated as the inverse of the first ISI (1/ms) [inv_first_ISI] |  |  |
| Instantaneous firing rate calculated as the inverse of the second ISI [inv_second_ISI:] |  |  |
| Instantaneous firing rate calculated as the inverse of the last ISI [inv_last_ISI] |  |  |
| Average difference between consecutive ISIs normalized to their sum [adaptation_index2:] |  |  |
| Voltage baseline before current step [voltage_base] |  |  |
| Voltage baseline after current step [voltage_after_stim] |  |  |
| Spontaneous AP count before current step [AP_count_before_stim] |  |  |
| Spontaneous AP count after current step [AP_count_after_stim] |  |  |
| Difference between maximum and minimum AP amplitudes [max_amp_difference] |  |  |
| Ratio between AP counts evoked within the first and second half of the current step clustering_index: |  |  |
| Avg. fast AHP depth [fast_AHP:] |  |  |
| **Protocol 3** | **All Conditions: Step(s): -200 pA** |  |
| 1st AP amplitude [AP1_amp_rev] |  |  |
| 2nd AP amplitude [AP2_amp_rev:] |  |  |
| time_to_first_spike |  |  |
| inv_first_ISI |  |  |
| inv_second_ISI |  |  |
| inv_last_ISI |  |  |
| voltage_base |  |  |
| voltage_after_stim |  |  |
| AP_count_before_stim |  |  |
| AP_count_after_stim |  |  |
| Relative difference (in percentage) between voltage steady state and voltage deflection, both measured as the difference from baseline [sag_amplitude] |  |  |
| voltage_deflection:<br>Difference between minimum voltage value and baseline |  |  |
| **Protocol 4** | Membrane time constant [decay_time_constant_after_stim2:] |  |
| **Protocol 5** | Ratio between voltage deflection and current amplitude [input_resistance] |  |

### Fitting neuron models

We define a TC model comprising 26 free parameters, including the conductance densities along with the decay time constants of intracellular Ca^2+^ concentration (see *“Intrinsic membrane properties”*). We also added several voltage shift variables and multiplicative factors as free parameters to the equations describing the dynamics of Na_T_, K_DR_. K_M_. K_M_, and BK currents, with different effects on the firing properties generated by our models. Specifically, we added:

- Multiplicative factors to the equations of the rates for activation and inactivation of Na_T_ and K_DR_, fixing the ranges in a way that slowed down their dynamics, and thus regulated the AP width.
- A multiplicative factor to the equation of the rates for the activation of K_M_, fixing the ranges in a way that slows down the de-activation, and thus calibrated the contribution to firing rate adaptation.
- A voltage shift variable to the equations of activation and inactivation, along with their rates, for K_D_, fixing the range in a way that increased the impact of the channel on the first-spike latency, yielding half-values of activation and inactivation more similar to the values observed in pyramidal cells (Storm, 1988).
- Multiplicative factors and shift variables to the equations describing activation and inactivation, along with their time constants, for BK, mimicking the effects of β subunits expression on the channel dynamics (Behrens *et al*., 2000; Contreras *et al*., 2012), to modulate the impact of this channel on firing rate adaptation.

The values of the free parameters were determined by multi-objective optimization, using the BluePyOpt toolkit (Van Geit *et al*., 2016) to fit model traces to a set of target features (Table 2). These features described membrane potential dynamics (e.g., action potential amplitudes, after-hyperpolarization depth, spike count, firing adaptation index, and so on) in response to current injection paradigms and were extracted from our published whole-cell recordings in normal and parkinsonian states (Bichler *et al*., 2021). For each condition, model candidates were obtained with 15 optimization sessions, using 15 different random seeds, with 100 individuals and 100 generations per session. Technically, the optimizer minimized the error associated with each model as measured by the difference between electrophysiological features from simulations and experimental traces. The overall fitness of a given model resulted from the sum of the absolute errors associated with feature differences (passive features and active features, Table 2), each calculated as the deviation from the experimental mean normalized to the experimental standard deviation. To measure the firing properties of each model, the optimizer ran a battery of simulations reproducing experimental traces (Table 2). For final model selection, we first made a hall-of-fame as the population of models for which all errors fell within 3 standard deviations from mean (normal: n=686; 6-OHDA: n=4900). Of these models, we retained the ones that passed several additional quality checks. First, we compared the entire f-I curves, obtained with protocol 1, along with the rebound spike count versus current relationships, obtained with protocols 2 and 3, retaining the hall-of-fames that yieldedspike counts below 3 standard deviations from the experimental mean throughout the entire range of stimulation (normal: n=78; 6-OHDA: n=922). Second, we selected the models that replicated the effects of XE991 application (10-20 μM) (Bichler *et al*., 2021), simulated by decreasing the M-type conductance density by 70% (Yeung and Greenwood, 2005). In the experiments, XE991 application in normal condition decreased the rheobase by ∼80 pA and shifted the f-I curves to the left, while the application did not alter the response in parkinsonian condition. For each model, we then compared the f-I curves with and without XE991, retaining normal and parkinsonian models generating distinguishable (Wilcoxon, p < 0.1; with shift in rheobase < 100 pA) and indistinguishable (Wilcoxon, p ≥ 0.9; with no shift in rheobase) curves (normal: n=39; 6-OHDA: n=21). Third, by inspecting traces, we rejected a few models that generated a non-physiological after-hyperpolarization, with peaks below baseline (normal: n=28; 6-OHDA: n=19). Fourth, we tested the models with realistic synaptic inputs (see *Modeling synaptic inputs*), retaining the models that generated firing rates between 2.5 Hz and 90 Hz (normal: n=17; 6-OHDA: n=12).

### Modeling synaptic inputs

We modeled four groups of synaptic inputs: glutamatergic drivers (DRI), glutamatergic modulators (MOD), gabaergic inputs from substantia nigra reticulata (SNR), and gabaergic inputs from reticular thalamic nuclei (RTN), each defined by postsynaptic location, numbers of terminals, and unitary conductance (Table 3), representing the presynaptic activity by artificial spike trains with realistic firing rates and irregularity (i.e., coefficient of variation of inter-spike interval).

**Table 3.** Estimates of unitary synaptic conductance for each synaptic input to ventromedial motor thalamus. Summary of estimated unitary conductance for each group of synaptic inputs to ventromedial motor thalamus. Abbreviations: ***MOD,*** modulators; ***RTN,*** reticular thalamic nuclei; ***SNR,*** substantia nigra reticulata; ***DRI,*** drivers. References: ***1,*** (Laudes *et al*., 2012); ***2,***; ***3,***(Gornati *et al*., 2018b). Conductance values yielded amplitudes of inhibitory (IPSC) and excitatory (EPSC) postsynaptic currents (±SE) that match the amplitude measured in patch-clamp experiments on ventrobasal (Laudes *et al*., 2012) and ventromedial (Edgerton and Jaeger, 2014a; Gornati *et al*., 2018b) thalamic nuclei. In one experiment (Laudes *et al*., 2012), one reported the amplitudes for miniature IPSCs and EPSCs.

|  | Simulation | Experiment | Ref. |
| --- | --- | --- | --- |
| **MOD** | 2.2 nS<br>mEPSC: $-28.6 \pm 0.2$ pA | mEPSC: $-28.4 \pm 2.1$ pA | 1 |
| **RTN** | 0.8 nS<br>mIPSC: $26.6 \pm 0.6$ pA | mIPSC: $24.43 \pm 0.75$ pA | |
| **SNR** | 1.3 nS<br>mIPSC: $10.9 \pm 0.3$ pA with 1 active terminal | Minimum IPSC: 11.1 pA | 2 |
| **DRI** | 1.7 nS yielding $162.3 \pm 0.8$ pA with 4 active terminals | EPSC: $-165.0 \pm 40.2$ pA with ~4 active terminals | 3 |

**Table 4.** Circular statistics of spike-phase locking in normal and parkinsonian stats with oscillatory synaptic inputs.

| **Modulated inputs (s)** | **Beta band** | **Case** | **mean** | **SD** |
| --- | --- | --- | --- | --- |
| **SNR** | **12.5** | **+SNR, 6-OHDA** | 249.3 | 152.5 |
|  |  | **6-OHDA** | 265.9 | 161.0 |
|  |  | **Control** | 240.6 | 162.3 |
|  | **25** | **+SNR, 6-OHDA** | 313.2 | 155.4 |
|  |  | **6-OHDA** | 325.4 | 160.7 |
|  |  | **Control** | 313.0 | 161.5 |
| **SNR+MOD** | **12.5** | **+SNR, 6-OHDA** | 267.5 | 163.1 |
|  |  | **6-OHDA** | 283.7 | 174.0 |
|  |  | **Control** | 263.9 | 182.2 |
|  | **25** | **+SNR, 6-OHDA** | 301.7 | 156.7 |
|  |  | **6-OHDA** | 317.4 | 164.3 |
|  |  | **Control** | 294.9 | 169.7 |
| **SNR+MOD+DRI** | **12.5** | **+SNR, 6-OHDA** | 251.2 | 127.4 |
|  |  | **6-OHDA** | 264.5 | 131.6 |
|  |  | **Control** | 235.1 | 134.4 |
|  | **25** | **+SNR, 6-OHDA** | 237.6 | 131.4 |
|  |  | **6-OHDA** | 258.3 | 144.0 |
|  |  | **Control** | 217.5 | 131.4 |

Specifically, excitatory synapses comprised AMPA and NMDA components (Cavarretta *et al*., 2018), with a reversal potential of 0 mV, and NMDA/AMPA ratios of 0.6 for drivers and 1.91 for modulators as estimated from ventrobasal thalamus (Miyata and Imoto, 2006). For inhibitory synapses, we set the decay time constant to 14 ms (at 32° C) (Edgerton and Jaeger, 2014b), reversal potential to –76.4 mV (cf −75.6 mV) (Pathak *et al*., 2007), and Q_10_ to 2.1 (Otis and Mody, 1992).

For each group of synapses, we determined subcellular distributions and numbers of contacts. We fixed the proportion between DRI and MOD terminals to 10%, consistent with previous anatomical studies (Godwin *et al*., 1996; Van Horn *et al*., 2000; Van Horn and Sherman, 2007). The DRI terminals were located at a distance from soma that reproduced the distribution observed for deep cerebellar nuclei terminals in VM (Gornati *et al*., 2018a). For the other synaptic inputs, we accounted for the data obtained by electron microscopy of mouse VM (Dr. Yoland Smith, unpublished data), which yielded estimates of the linear density on transversal sections of dendrites for round-small terminals, small symmetric terminals, and large symmetric terminals, putatively MOD, RTN, and SNR terminals, respectively, grouped in three brackets on the dendritic diameter (0-0.5 μm, 0.5-1 μm, ≥ 1 μm). For each, we assumed that the distance between consecutive terminals along the longitudinal axis of the dendrites was equal to the diameters of terminals, which is ∼0.8 μm for MOD and RTN terminals (Dr. Yoland Smith, unpublished data), and 2.8 μm for SNR terminals (Bodor *et al*., 2008). However, our electron microscopy data showed the distributions of terminals on dendrites only. Therefore, in our simulations, we added the SNR terminals contacting the TC soma, which accounted for 10% of total SNR synapses (Bodor *et al*., 2008).

After defining the subcellular distributions, we estimated the unitary synaptic conductance for each group of synapses. To this end, we designed an optimizer that determined the optimal values matching the postsynaptic current amplitudes observed experimentally (Table 3, Experiment). The optimizer explored the range of conductance values by using a convergence algorithm, testing each value by running simulations that reproduced the patch-clamp experiments (Table 3, Simulation).

By applying the subcellular distributions of synaptic terminals described above to the morphology (AA0719; MouseLight Archive), we obtained estimates of the number of terminals for each group of synapses (MOD: 3625; DRI: 350; SNR: 25; RTN: 400). In our simulations, we decreased the MOD terminals to 1450, i.e., ∼40% of the estimated value, accounting for the proportion of silent and non-silent pyramidal neurons observed in layer VI of motor cortex of cats (Sirota *et al*., 2005). This configuration yielded a basal firing rate of 197.7±80.1 Hz (±SD) in normal conditions, higher than observed experimentally (∼20 Hz) (Inagaki *et al*., 2022), suggesting a non-physiological excitation-inhibition balance. As the sample size of large dendrites (≥ 1 μm in diameter) was the smallest in our electron microscopy experiment, we assumed that the number of SNR terminals was underestimated. We then tested the increase in the number of SNR terminals to 100, which yielded a firing rate of 25.5±16.3 Hz (±SD) in normal conditions, consistent with the values observed experimentally (Inagaki *et al*., 2022).

### Pre-synaptic activity: artificial spike train generation

To represent presynaptic activity, we generated artificial spike trains (Abbasi *et al*, 2020), with firing rates consistent with experimental estimates (DRI: 30 Hz; MOD: 1.1 Hz; RTN: 10 Hz; SNR: 50 Hz) (Sirota *et al*., 2005; Huh and Cho, 2016; Barrientos *et al*., 2019; Inagaki *et al*., 2022). Specifically, we used an algorithm that generates a random sequence of inter-spike intervals picked from a gamma distribution with a refractory period and “shape” parameter set a priori. We targeted generic spike properties with a coefficient of variation of inter-spike intervals (CV_ISI_) of 0.45 (as observed in SNR during in vivo recordings) (Lobb and Jaeger, 2015), obtained with a shape of 5 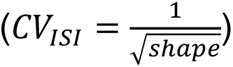 and a refractory period of 3 ms. Unless otherwise noted, we maintained these combinations of parameters for all the synaptic inputs (see Figs. 4 & 5). We enhanced the published method to generate bursts, treated as a special case of event in each spike train. Specifically, each burst sequence was generated separately and subsequently merged with an existing baseline firing spike train. A template described the time course of the intra-burst firing rate, which can be manipulated to fit different accelerando-decelerando patterns. Specifically, we used different values of shape for the gamma distributions of inter-spike intervals for regular spike trains (=5) and bursts (=3), which yielded bell shaped and more symmetric distributions for the former, and highly right skewed ones for the latter (Selinger *et al*., 2007). Furthermore, inter-burst intervals were governed by a gamma distribution, associated with mean and regularity parameters that correlates with average (positively) and variance (negatively) of the inter-burst intervals, respectively.

**Figure 4.**
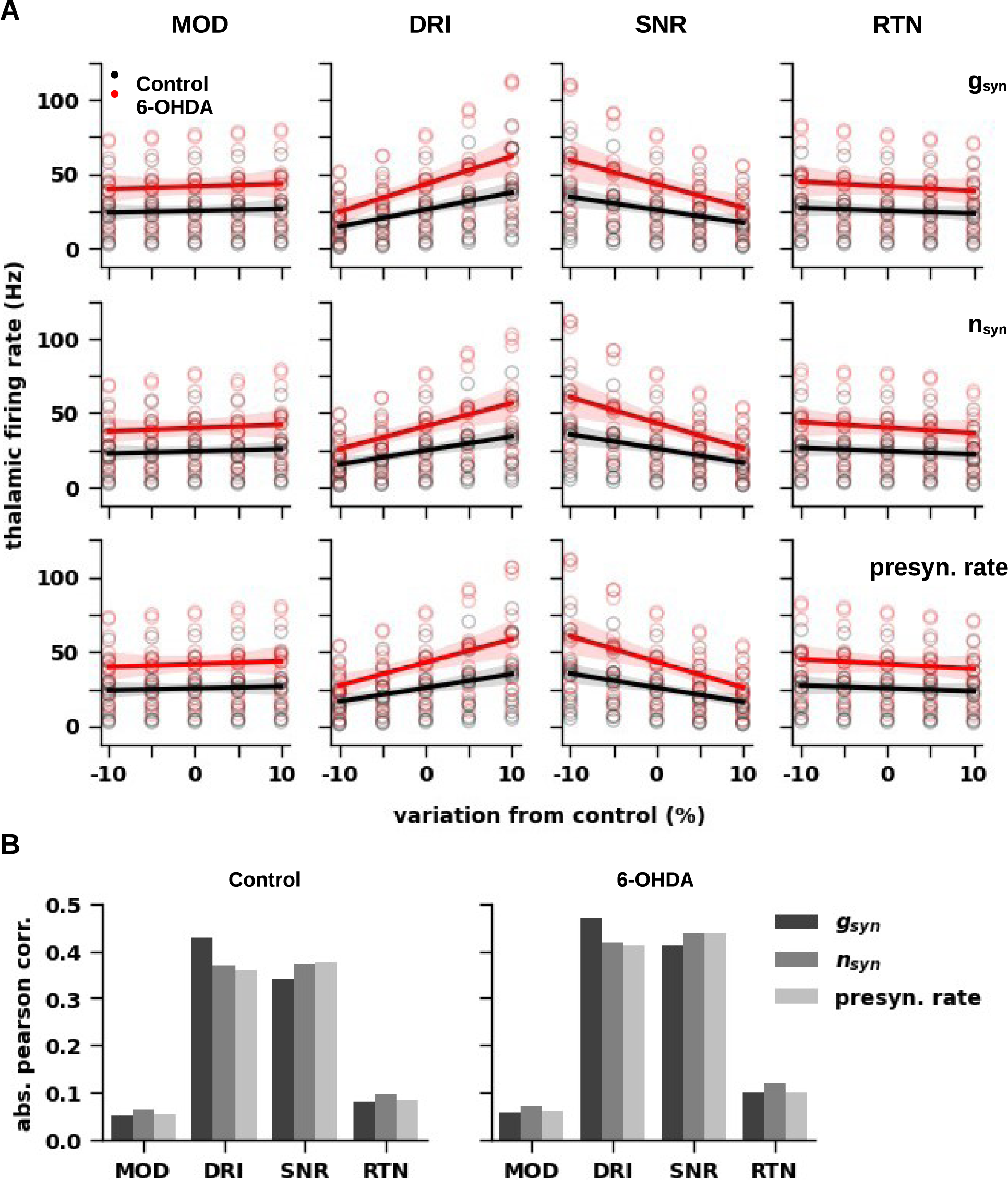
Sensitivity analysis of synaptic inputs. ***A,*** Dependence of the firing rate of thalamocortical neurons in normal (black) and parkinsonian (red) states on parameter variations (±10% from control) for synaptic inputs, i.e., individual synaptic conductance (g_syn_), number of synapses (n_syn_), and presynaptic firing rate, for modulators (MOD), drivers (DRI), substantia nigra reticulata (SNR) and reticular thalamic nuclei (RTN). ***B,*** Linear (Pearson) correlation of thalamic firing rates with the percentages of variation from control for each parameter indicated in *A*. Variations in the parameters of DRIs and SNR yielded the most significant linear correlations (p < 0.001).

**Figure 5.**
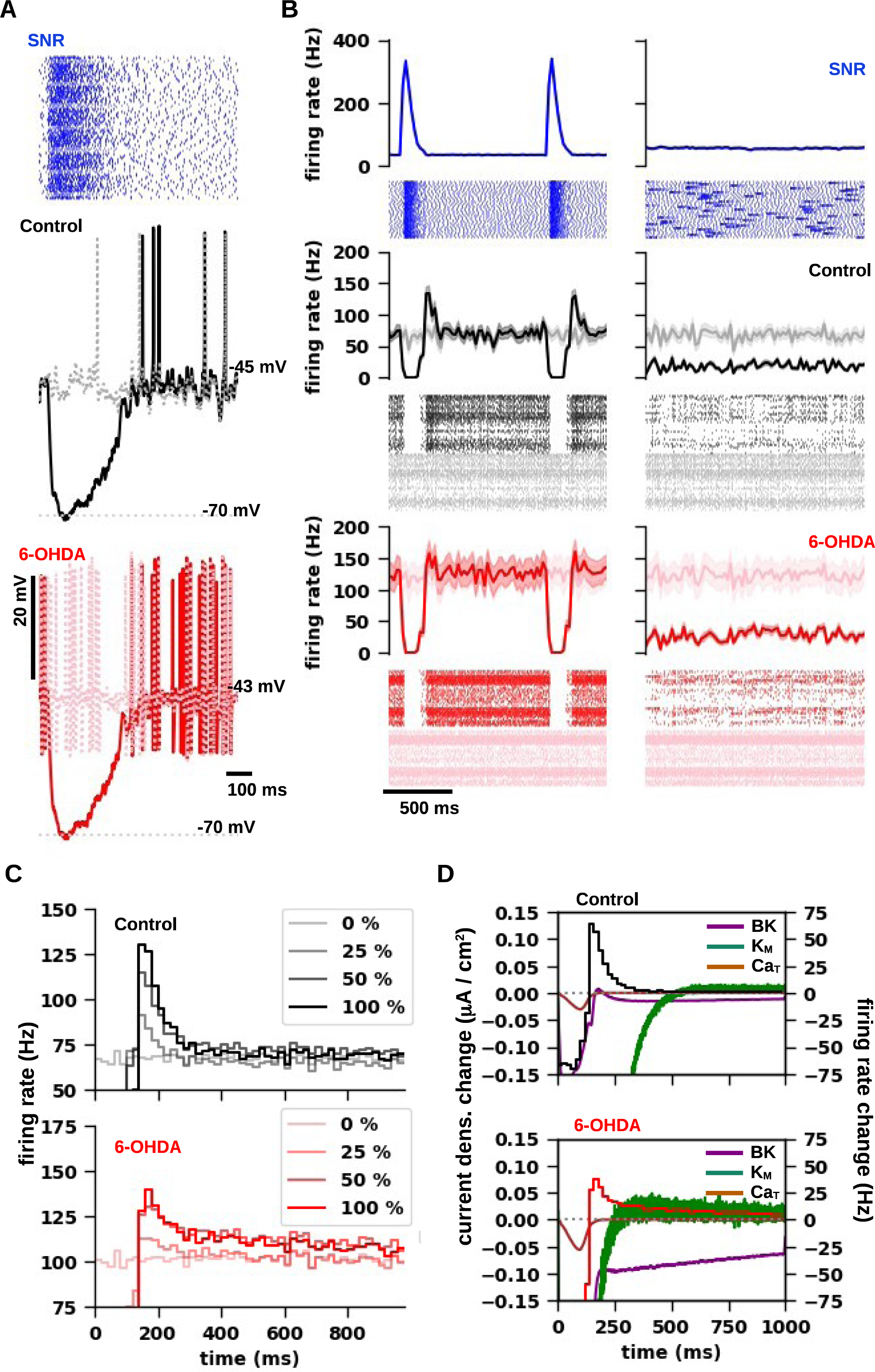
Synchronous bursting in substantia nigra reticulata evokes rebound activity in motor thalamocortical neurons. ***A,*** Representative responses of thalamocortical neuron models in normal (Control; middle, black) and parkinsonian (6-OHDA; bottom, red) states to inputs from substantia nigra reticulata (SNR) with synchronous bursting (top, blue). The gray dashed traces correspond to the responses in normal (top) and parkinsonian (bottom) states without bursting in SNR (not shown). ***B,*** Thalamocortical neuron responses in normal (black) and parkinsonian (red) states to SNR inputs (blue) with synchronous (right) and asynchronous (left) rebound bursting. ***Top,*** Spike histograms and pooled raster plots of the activity from SNR for 10 simulations with synchronous bursting. Rebound bursting was ∼150 ms in duration, with an average intra-burst rate of ∼170 Hz, and an inter-burst rate of 35 Hz, yielding a firing rate of 48.3±0.4 Hz (±SD) with a coefficient of variation of inter-spike interval of 0.57±0.01. Above the rastergrams, the curves show instantaneous firing rate versus time for SNR activity with (blue) and without (gray) bursting. ***Middle, Bottom,*** The corresponding spike histograms and raster plots of thalamic activity in presence of bursting in SNR in normal (black) and parkinsonian (red) states, or without bursting in SNR (gray). Above the rastergrams, the curves show instantaneous firing rate versus time for the activity of thalamocortical neurons with (red, black) and without (gray) bursting in SNR. ***C,*** Post-stimulus spike time histograms (PSTH) of thalamocortical neuron activity in normal (Control; top, red) and parkinsonian (6-OHDA; bottom, red) conditions, with different percentages of SNR inputs generating synchronous bursting (100%, 50%, 25%, 0%). The PSTHs were averaged for different models (Control: n=17; 6-OHDA: n=9) and 10 trials, and over time window of 1000 ms (1 second) in length, starting from the SNR bursting onset. ***D,*** Changes in density of M-type potassium (green), BK potassium (purple), and T-type Ca^2+^ (brown) currents during post-inhibitory firing activity in normal (Control; top) and parkinsonian (6-OHDA; bottom) models of thalamocortical cells with 100% of SNR inputs generating synchronous bursting. For each current, the changes were calculated as the difference between the curves obtained with 100% and 0% of SNR inputs generating synchronous bursting. The background of each panel shows the difference between PSTHs obtained with 100% and 0% of synchronous SNR bursting for normal (top) and parkinsonian (bottom) models.

We then generated bursting activity for SNR inputs (see Fig 5), as artificial spike trains of ∼150 ms in duration, with an average firing rate of ∼170 Hz and a refractory period of 1.5 ms. For the inter-burst activity, we decreased the baseline firing rate to 35 Hz and increased the regularity to 50. The inter-burst intervals were 1s on average, with a minimum value of 150 ms and regularity values of 5 and 50000 for spike trains with asynchronous and synchronous bursting, respectively. Overall, this configuration yielded artificial spike trains with firing rate of 48.5±1.0 Hz (±SD) and coefficient of variation of inter-spike interval of 0.57±0.02, consistent with our experimental recordings from SNR (Lobb and Jaeger, 2015).

### Data analysis

We used a custom version of the eFEL python package for feature extraction from membrane potential traces (https://efel.readthedocs.io). The junction potential of the intracellular medium was estimated using JPcalc software (Barry, 1994). For each simulated experiment, reversal potentials of ion species were estimated by using the Nernst equation, accounting for the composition of the aCSF (see Table 1). For statistical analyses, we used Python3 with the SciPy and NumPy packages. To compare f-I curves (Fig. 2), we used the Wilcoxon matched-pairs signed rank test. To compare different sets of measures, we used Mann-Whitney U tests. To assess the significance of the Pearson correlation (Fig. 3), we used the Kowalski test. To assess the significance of spike-phase locking (Fig. 6, 7), we used the Rayleigh test on polar data.

**Figure 6.**
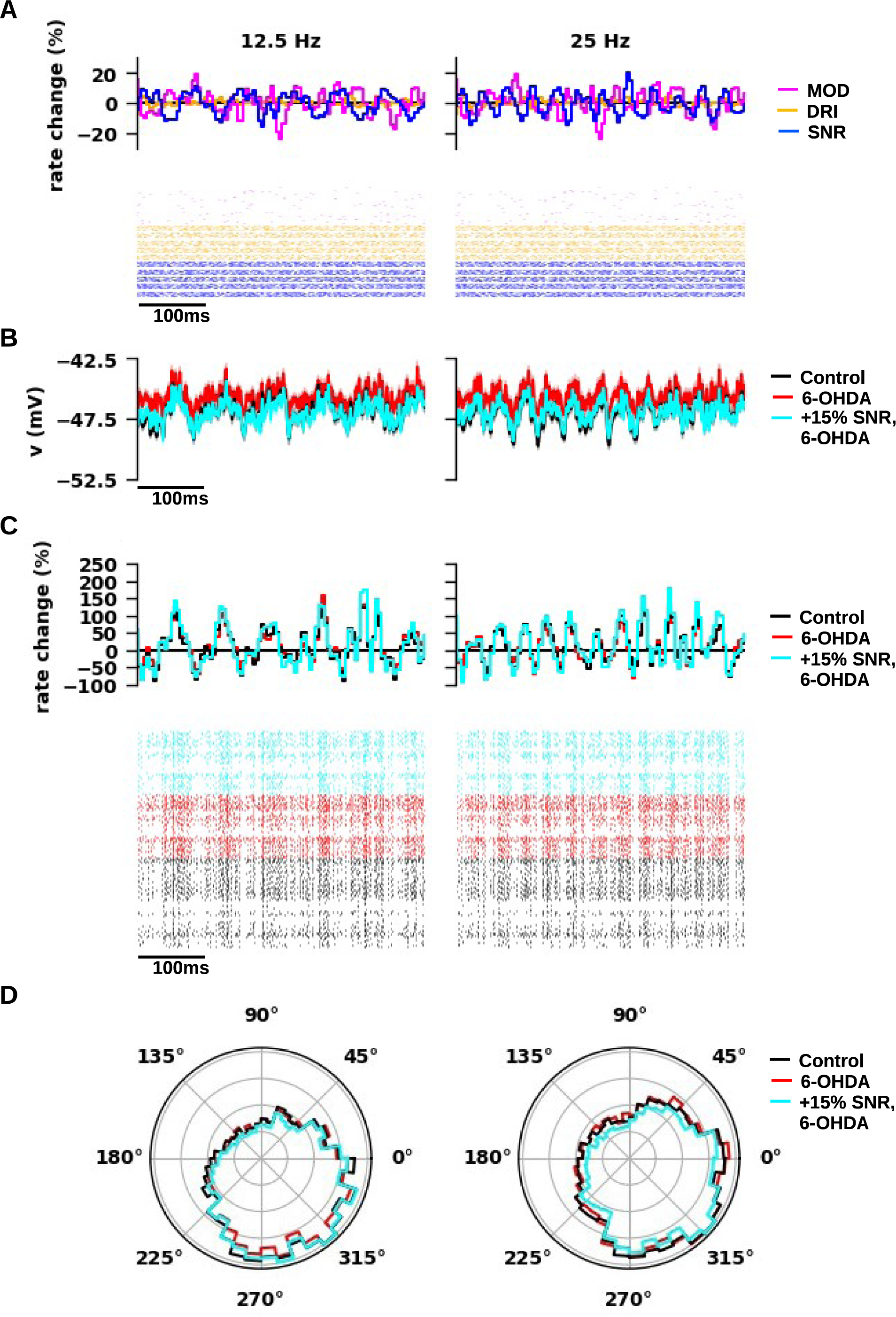
Beta modulation of inhibitory inputs from substantia nigra reticulata. ***A,*** Beta modulation in the activity of substantia nigra reticulata (SNR) at 12.5Hz (left) and 25Hz (right). Relative variation of instantaneous firing rate to average versus time(Top) and raster plots (Bottom) of the presynaptic activity for MOD (magenta), DRI (orange), and SNR (blue) for different seeds (n=10). The rastergrams of each input were a subset of total (*n*=100). ***B,*** Somatic voltage traces of thalamocortical neurons in normal (black) and parkinsonian (6-OHDA) states with regular (red) and increased (+15%; cyan) SNR conductance for parkinsonian models. ***C,*** Relative variation of instantaneous firing rate to average versus time(Top) and rastergrams (Bottom) of thalamocortical neuron activity in normal (black) and parkinsonian states with regular (red) and increased (+15%; cyan) SNR conductance for parkinsonian models. ***D,*** Phase plots of the spiking activity shown in C for normal and parkinsonian models of thalamocortical neurons (Rayleigh, p < 0.01).

**Figure 7.**
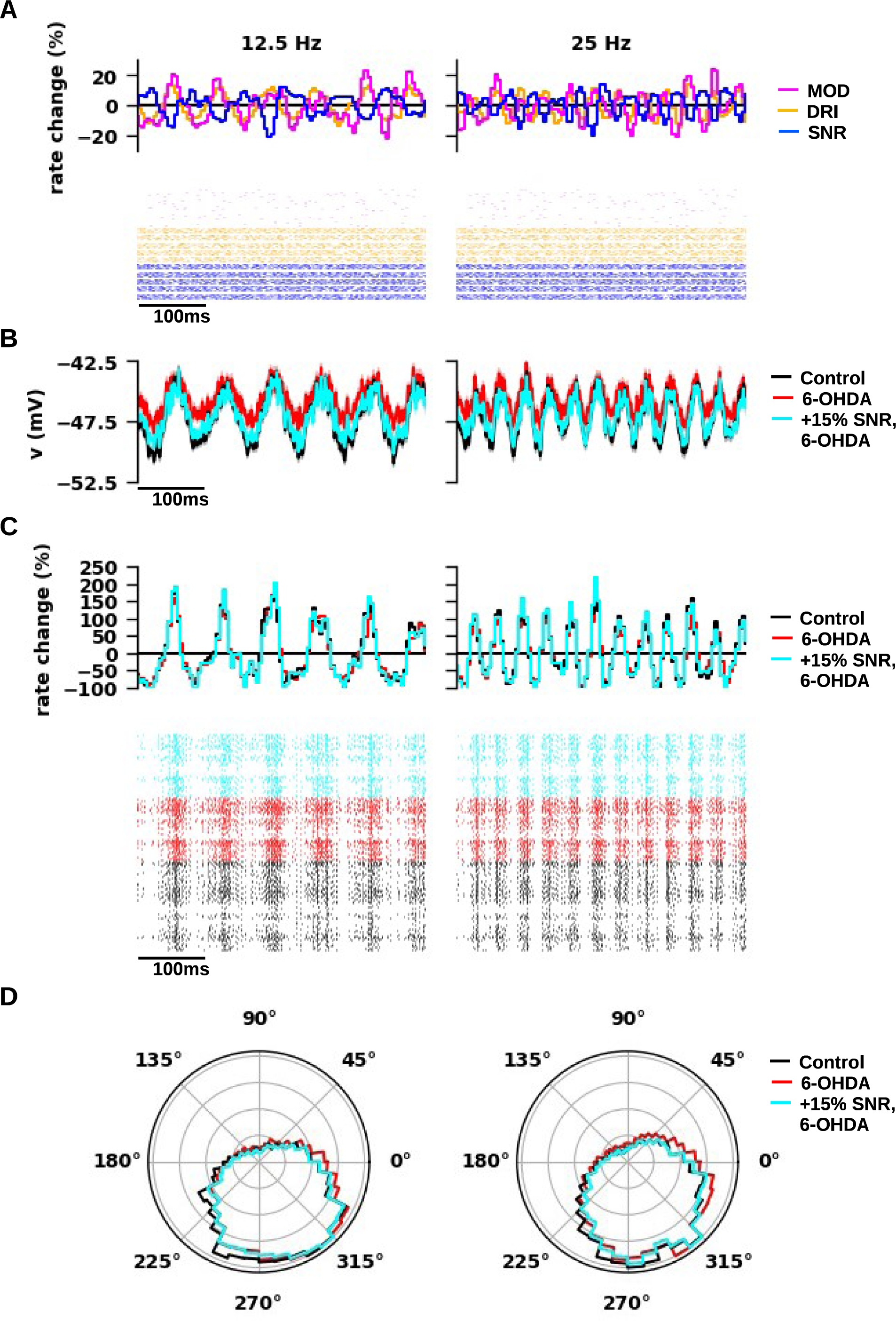
Beta modulation of inhibitory inputs from substantia nigra reticulata and excitatory modulators and drivers. ***A,*** Beta modulation in the activity of substantia nigra reticulata (SNR), modulators (MOD), and drivers (DRI), at 12.5Hz (left) and 25Hz (right). The oscillations in MODs and DRIs were shifted by 180^∘^ with respect to SNR inputs. Relative variation of instantaneous firing rate to average versus time (Top) and raster plots (Bottom) of the presynaptic activity for MOD (magenta), DRI (orange), and SNR (blue) for different seeds (*n*=10). The rastergrams of each input were a subset of total (*n*=100). ***B,*** Somatic voltage traces of thalamocortical neurons in normal (black) and parkinsonian (6-OHDA) states with regular (red) and increased (+15%; cyan) SNR conductance parkinsonian models. ***C,*** Relative variation of instantaneous firing rate to average versus time(Top) and rastergrams (Bottom)of thalamocortical neuron activity in normal (black) and parkinsonian states with regular (red) and increased (+15%; cyan) SNR conductance for parkinsonian models. ***D,*** Phase plots of the spiking activity shown in *C* for normal and parkinsonian models of thalamocortical neurons (Rayleigh, p<0.001).

### Results

In our previous study, we performed *in vitro* experiments to investigate the effects of dopamine depletion on ventromedial thalamus (VM) of mice, induced by unilateral 6-OHDA lesions (Bichler *et al*., 2021). We found that the dopaminergic loss enhanced the excitability of thalamocortical neurons (TC), because of M-type potassium current suppression. To understand how these excitability changes may impact synaptic integration in vivo, we constructed biophysically detailed VM TC models in normal and 6-OHDA depleted conditions (Fig. 1; for details, see Method section *“Fitting neuron models“*), using the NEURON simulator (Hines and Carnevale, 1997). We then modeled the main afferent inputs to VM, i.e., excitatory modulators (MOD), excitatory drivers (DRI), inhibitory inputs from substantia nigra reticulata (SNR), and inhibitory inputs from reticular thalamic nuclei (RTN), to simulate *in vivo* conditions (Fig. 1; for details, see Method section *“Simulation of in vivo conditions“*). This approach allowed us to assess the interaction of intrinsic thalamic neural dynamics with synaptic input properties expected in normal or parkinsonian waking conditions through measurements in the model that cannot be performed experimentally in vivo.

### Simulation of in vitro experiments

Physiologically, VM TCs display the characteristic firing modes, e.g., tonic firing, low-threshold spiking, and rebound bursting (Edgerton and Jaeger, 2014b; Bichler *et al*., 2021), commonly observed in other thalamic nuclei (Llinás and Steriade, 2006), while dopamine depletion increases tonic firing and prolongs rebound bursting in these neurons (Bichler *et al*., 2021). Here we tested the ability of the VM TC models in reproducing *in vitro* recordings and the changes following dopamine depletion, comparing simulation and experimental responses.

Specifically, we considered whole-cell recordings from slices of adult mice VM in normal conditions and after unilateral 6-OHDA application (control: n=9; 6-OHDA: n=7), stimulated by depolarizing (20-300 pA; 2 s of duration; Fig. 2 A, B) or hyperpolarizing (from –50 pA to –200 pA; 2 s of duration; Fig. 2C, D) current steps on top of a bias current (control: 93.3±18.7; 6-OHDA: 67.1±12.7; ±SE). In particular, the bias current held the membrane potential to –69 mV, inactivating T-type Ca^2+^ channels, and thus enabled the generation of tonic firing in response to depolarizing steps (Fig. 2 A, left), while hyperpolarizing steps evoked rebound bursting (Fig. 2 C, left). 6-OHDA enhanced the excitability of thalamocortical neurons, increasing responses of TCs in tonic firing (Fig. 2A, left; compare black and red traces), assessed as a shift of the f-I curves to the left (Wilcoxon, p < 0.01; Fig. 2B, left; compare black and red curves), and prolonged rebound bursting (Fig. 2A, left; compare black and red traces), assessed as an increase in the rebound spike count (Mann-Whitney, 20-40 mV: p=0.015; 40-60 mV: p < 0.01; Fig. 2D, left; compare black and red bars for each bracket).

To allow a direct comparison between the two sets of traces, the simulations reproduced the stimulation protocols (e.g., step and bias current amplitudes) and the environmental conditions of slices (e.g., temperature, reversal potential of ion species, holding membrane potential; Table 1). We thus observed that the models (control: n=17; 6-OHDA: n=12) replicated the responses in tonic and burst firing modes observed experimentally, in response to depolarizing and hyperpolarizing steps, respectively, with realistic levels of firing rate adaptation and action potential properties (e.g., AP height, AHP depth, firing rate adaption). Compared to normal models, parkinsonian models displayed stronger excitability, generating more action potentials in response to depolarizing and hyperpolarizing current steps (Fig. 2 A, C, right; compare black and red traces). Parkinsonian models generated significantly higher spike rates throughout the entire range of stimulation using depolarizing currents (Wilcoxon, p < 0.01; Fig. 2B, right; compare black and red curves), while the spike rates were comparable to the values observed experimentally for both normal and parkinsonian models (Fig. 2B; compare experimental and simulation f-I curves in left and right panels, respectively). Likewise, rebound spike counts were statistically different in normal and parkinsonian models (Mann-Whitney, 20-40 mV: p < 0.01; 40-60 mV: p < 0.01; Fig. 2D, compare black and red bars for each bracket) but comparable to the corresponding experimental values (Fig. 2D; compare experimental and simulation spike counts for each bracket in left and right panels, respectively).

The models discussed above resulted from a selection process consisting of several quality checks. In particular, one check concerned the ability of the models in replicating the effects of XE-991 application (10-20 μM) (Bichler *et al*., 2021), simulated by reducing the M-type conductance density by 70% (Yeung and Greenwood, 2005), which shifted the f-I curves to the left significantly for normal but not parkinsonian models (not shown), in accord with our findings *in vitro* (Bichler *et al*., 2021). This yielded parkinsonian models with significantly lower expression of M-type potassium current than control (Mann-Whitney, p < 0.01; control: 314.2±117.3 μS/cm^2^; 6-OHDA: 117.1±10.6 μS/cm^2^). Increasing M-type potassium current in parkinsonian models by ∼32 times rescued the effects of dopamine depletion (not shown), making the f-I curves of parkinsonian models indistinguishable from control (Wilcoxon, p=0.73), and reducing their rebound spike count to similar values as observed for control (Mann-Whitney, 20-40 mV: p=0.02; 40-60 mV: p=0.13). Taken together, these simulations demonstrated that our models replicate the firing behaviors of VM TCs observed *in vitro* along with the alterations caused by dopamine depletion.

### Simulation of in vivo conditions

After fitting the TC models to in vitro recordings, we modeled the main afferent inputs to VM (Fig. 3): 1) modulators (MOD), approximating the glutamatergic inputs conveyed from layer VI of Motor Cortex (MC); 2) drivers (DRI), approximating glutamatergic inputs from layer V of MC and motor-related subcortical areas; gabaergic inputs from 3) substantia nigra reticulata (SNR) and 4) reticular thalamic nucleus (RTN).

For each group of synapses, we constrained synaptic physiology, subcellular distributions, number of terminals, and individual conductance to experimental data (for detail, see Method section *“Modeling synaptic inputs”*). In particular, the distribution of DRI terminals reproduced the distance from soma observed for deep cerebellar nuclei terminals in VM (Fig. 3 A, B) (Gornati *et al*., 2018a), while the distributions for MODs, RTN, and SNR were modeled on our electron microscopy data (Fig. 3 A, B; for detail, see Method section *“Modeling synaptic inputs”*). As an emergent property, we obtained a co-localization along the somato-dendritic arbor of VM TCs (Fig. 3B) for DRI and SNR terminals, targeting proximal dendrites with larger diameters, as well as MOD and RTN terminals, targeting distal dendrites with small diameters, suggesting that a stronger interaction take place in VM TCs between each pair of co-localized terminals.

We then tested the impact of these synaptic inputs on normal and parkinsonian TC models (Fig. 3C). We represented the pre-synaptic activity with artificial spike trains replicating firing rates and irregularity observed experimentally (Fig. 3Ca; for detail, see Method section *“Modeling synaptic inputs”*). With these configurations of synaptic inputs, we observed that the highest values of total synaptic conductance over time were achieved by SNR inputs and DRIs (Fig. 3Cb), suggesting that these two classes of synaptic inputs sustained the TC firing in normal and parkinsonian states (see also *“Sensitivity analysis of synaptic response function”*). Although the two populations of models received the same synaptic inputs, they generated different firing responses (Fig. 3Cc). Compared to control, parkinsonian models displayed increased firing rates (Mann-Whitney, p < 0.0001; normal: 25.5±16.3 Hz; 6-OHDA: 41.8±22.4 Hz; ±SD) and baseline voltage (Mann-Whitney, p < 0.0001; normal: −46.8±1.4 mV; 6-OHDA: −45.5±1.3 mV). Therefore, the models suggest that the homeostatic changes underlying the increased excitability observed in vitro for parkinsonian states resulted in a strong firing increase with a balance of excitatory and inhibitory synaptic inputs as well.

### Sensitivity analysis of synaptic response function

To establish the importance of each synaptic afferent, we performed a sensitivity analysis of their parameters, i.e., numbers of synapses (*n_syn_*), unitary conductance (*g_syn_*), and presynaptic firing rate. For each group of synapses, we plotted the thalamic firing rate for normal and parkinsonian models as a function of each parameter, varied by ±10% from control (Fig. 4A). For each parameter, we performed a linear regression between the percent of variation and the thalamic firing rate in normal and parkinsonian states (Fig. 4A, black and red lines), and used the absolute value of the Pearson correlation between thalamic firing rate and the varying parameter (Fig. 4B).

In normal and parkinsonian states, increasing synapse number, conductance, or firing rate for RTN and SNR inputs decreased firing rates of the TC models (i.e., negative correlation), while increasing these parameters for MODs and DRIs resulted in the opposite effect (i.e., positive correlation; Fig. 4A). The fitted lines for each parameter were steeper for DRI and SNR, whereas they appeared flat for MODs and RTN (Fig. 4A), and the firing rate was higher in the parkinsonian state than the control state throughout the entire range of variation. By comparing the absolute values of correlation, we observed strong (0.34-0.47; Fig. 4B) and significant correlations for all the parameters related to DRIs and SNR (Kowalski test, p < 0.0001), whereas weak correlations were observed for MODs and RTN (0.05-0.12; Fig. 4B), insignificant for all the parameters related to MODs and RTN except the number of RTN synapses, which however displayed a weaker significance than DRIs and SNR (Kowalski test, normal: p=0.005; 6-OHDA: p=0.004). Taken together, these simulations suggest that, under these configurations of afferent inputs, VM output *in vivo* is primarily driven by DRIs and SNR but not MODs and RTN, without any relevant difference between normal and parkinsonian states. This is consistent with the observation that synaptic inputs from SNR and DRIs yielded the highest values of total synaptic conductance over time (see Fig. 3). Additionally, synaptic inputs evoked stronger responses in parkinsonian states throughout the entire range of parameters, suggesting a robust effect of dopamine depletion on VM TC excitability.

### Effects of synchronous bursting inputs from substantia nigra reticulata

Numerous speculations have described rebound bursting as a hallmark of parkinsonian pathophysiology (for review, see (Llinás and Steriade, 2006)), while experiments showed that the occurrence of rebound bursting in ventrolateral thalamus coincides with the emergence of motor deficits (Kim *et al*., 2017). *In vitro* experiments showed that synchronous activation of nigral axons could evoke rebound bursting in VM, where dopamine depletion enhanced rebound bursting evoked by hyperpolarizing current steps (Bichler *et al*., 2021). Additionally, *in vivo* experiments showed that dopamine depletion increased spike synchrony and bursting in SNR (Wang *et al*., 2010; Anderson *et al*., 2015; Willard *et al*., 2019), an ideal mechanism for evoking rebound spiking in VM. While these findings support the possibility that rebound spiking in VM might contribute to parkinsonian dynamics, this hypothesis has not yet been directly tested in behaving animals.

To investigate the effects of synchronous nigral bursting on VM output, we generated trains of nigral bursts reproducing the rates and duration observed in our experimental recordings in dopamine depleted mice (for detail, see Method section *“Pre-synaptic activity: artificial spike train generation”*). We observed that synchronous SNR bursts hyperpolarized the membrane of TCs (Fig. 5 A, B), blocking firing activity in normal (black) and parkinsonian (red) states, followed by a temporary firing rate increase at the offset of the nigral bursting. Comparing the thalamic activity evoked by SNR inputs with (Fig. 5A, red, blue) and without bursts (Fig. 5A, gray), we observed that the post-inhibitory increase in firing rate occurred robustly at the offset of the SNR bursting, without affecting the firing activity over different time windows (Fig. 5A, left). Instead, asynchronous SNR bursting decreased the average firing rates throughout the entire course of firing activity (Fig. 5B, right). By comparing the PSTHs with (black, red) and without (gray) synchronous nigral bursting (Fig. 5C), we observed that SNR bursting evoked a significant post-inhibitory firing rate increase in most TC models (normal: 17/17; 6-OHDA: 9/12), with different courses in normal and parkinsonian states, reaching a peak instantaneous firing rate of ∼125 Hz and ∼140 Hz, respectively, and lasting up to 380 ms and 760 ms, respectively. Both the firing rate peak and the duration of the post-inhibitory firing rate increase positively correlated with the percentage of synchronous SNR inputs, maintaining different peaks and durations in the two states. Therefore, our simulations suggest that synchronous SNR bursting evokes post-inhibitory spiking activity in motor thalamus with different courses in normal and parkinsonian states, and that the intensity of this effect correlates with the percentage of synchronous SNR inputs.

To seek a mechanistic explanation for the difference between the post-inhibitory responses observed in normal and parkinsonian models, we analyzed the variation in transmembrane current density induced by each HH-conductance over time (Fig. 5D). Surprisingly, the post-inhibitory increase in firing rate following synchronous SNR bursting was not sustained by T-type Ca^2+^ current, as the hyperpolarization was not sufficient to de-inactivate these channels (Fig. 5D, brown lines). Instead, the early increase at the offset of the hyperpolarization coincided with a state of incomplete recovery for M-type potassium current (Fig. 5A, green lines), observed with both with normal and parkinsonian models. Additionally, with parkinsonian models only, the incomplete recovery of BK potassium current, which maintained the current lower than baseline by ∼0.1 μS/cm^2^ for ∼800 ms, contributed to both early and late parts of post-inhibitory firing rate increase (Fig. 5D, bottom, purple line). Therefore, these predictions suggest that synchronous SNR bursting in vivo cannot de-inactivate T-type Ca^2+^ channels enough to evoke rebound bursting in TCs, while it causes a post-inhibitory increase in firing activity, due to the slow recovery of M-type potassium current for models in both states, along with the slow recovery of the BK potassium current in parkinsonian models only.

### Effects of beta modulation in the synaptic afferents to ventromedial thalamus

Another hallmark of parkinsonian pathophysiology is the exaggeration of beta oscillations in VM, observed by local field potential (LFP) recordings (Brazhnik et al., 2012; Brazhnik et al., 2016; Nakamura et al., 2021), with increased spike-LFP coherence in VM and SNR, MC (Brazhnik et al., 2012; Brazhnik et al., 2016; Nakamura et al., 2021). Therefore, it has been hypothesized that oscillations might originate in SNR and then entrain VM, propagating globally throughout the entire motor cortico-thalamocortical loop. We tested this hypothesis by introducing fluctuations in the firing activity of SNR within the beta band, i.e., at 12.5 Hz and 25 Hz (Fig. 6A, blue curve and raster), with amplitude ±10% as the baseline. We observed that somatic membrane potential (Fig. 6B) and instantaneous firing rate (Fig. 6C) reflected the fluctuations in SNR, with significant spike-phase locking (Rayleigh, p < 0.01; Fig. 6D). Compared to SNR, the oscillations in firing rate of VM TCs achieved higher amplitudes, with down and up phases of –100% and +150% circa, respectively, without any noticeable difference between normal and parkinsonian states, suggesting that VM does not only relay but also amplifies the beta oscillations in SNR in either state.

Moreover, canonical models of Parkinson Disease (PD) suggest that excessive nigrothalamic inhibition, due to increased SNR firing rate, might impede thalamocortical transmission of motor signals, preventing the regular course of movements (for review, see (DeLong, 1990)). While our simulations suggested that increased firing rate and/or synaptic conductance of SNR inputs could be two ways of enhancing nigrothalamic inhibition (see Fig. 3), studies on rodents led to inconsistent conclusions about the changes in SNR firing rate after dopamine depletion, with some experiments showing no changes or even a decrease in firing rate across different sets of SNR neurons (Díaz *et al*., 2003; Wang *et al*., 2010; Anderson *et al*., 2015; Lobb and Jaeger, 2015; Willard *et al*., 2019). Therefore, these findings corroborate the possibility that enhanced SNR inhibition could be a consequence of increased synaptic conductance rather than firing rate. We tested this hypothesis by increasing unitary synaptic conductance by 15% for SNR on parkinsonian models, yielding a basal firing rate indistinguishable from control (24.6±14.8 Hz vs 25.5±16.3 Hz; Mann-Whitney, p=0.47), consistent with previous observations from basal ganglia receiving nuclei of motor thalamus (Anderson *et al*., 2015; Di Giovanni *et al*., 2020; Nakamura *et al*., 2021). The increase did not affect the oscillatory firing patterns in the VM (Fig. 6A-D, compare magenta and dark cyan with dark pink and light cyan curves, respectively), suggesting that enhanced nigrothalamic inhibition did not increase spike-phase locking alone.

LFP recordings revealed that dopamine depletion increased beta oscillations in layer 5/6 of MC (Brazhnik *et al*., 2012) and the spike-phase locking with SNR (Brazhnik *et al*., 2012; Nakamura *et al*., 2021). To study how these oscillations could entrain VM, we added beta modulation in MODs and DRIs, approximating glutamatergic inputs to VM from layer 6 and 5 of VM in our simulations, respectively. Their phases were shifted by 180° with respect to SNR, consistent with previous experimental recordings (Brazhnik *et al*., 2012), which prevented a cancellation of inhibitory and excitatory input modulation. Compared to beta modulation in SNR alone, adding beta modulation in the MODs only did not affect the oscillatory firing activity in VM (not shown). Adding beta modulation in both the MODs and DRIs (Fig. 7A) yielded subthreshold oscillations in somatic voltage traces (Fig. 7B) as well as instantaneous firing rates (Fig. 7C) that were more demarcated than those observed with modulations of SNR alone (cf Fig. 6) or SNR and MODs (not shown), enhancing the significance of the spike-phase coherence (p < 0.0001). In particular, the oscillations in firing rate displayed down and up phases of – 100% and +200% circa, confirming that VM amplifies the beta oscillations in SNR and DRIs. Therefore, our simulations suggested that cortical inputs from layer 5 were more effective than those from layer 6 in transmitting beta oscillations to VM. Furthermore, increasing SNR conductance in parkinsonian models reduced the circular standard deviations of spike time by 15-20° when testing beta oscillations in MODs only (Table 3), becoming lower than control, whereas the effects were negligible with combined oscillations in MODs and DRIs, suggesting that the enhanced SNR inhibition refines the transmission of beta oscillations from SNR and MODs to VM.

### Discussion

We used biophysically detailed simulations to investigate the consequences of dopamine depletion on synaptic integration in VM (Fig. 1). We constructed two populations of VM TC models reproducing the firing properties observed in VM slices from control and 6-OHDA lesioned mice (Fig. 2) (Bichler *et al*., 2021). To simulate in vivo conditions, we modeled the main afferent inputs to VM (Fig. 3), i.e., glutamatergic MODs and DRIs, gabaergic inputs from SNR and RTN. Our simulations showed that the increased excitability observed in vitro after 6-OHDA lesioning (see Fig. 2) will also result in a strong firing rate increase in vivo (Fig. 3). The sensitivity analysis of the models of afferent inputs suggested that VM output *in vivo* is primarily driven by DRIs and SNR (Fig. 4). We then systematically tested the effects of pathological firing pattern changes observed in dopamine-depleted conditions: synchronous bursting (Fig. 5) and increased beta oscillations (Figs. 6 and 7). Synchronous SNR bursting evoked post-inhibitory spike rate increases in VM TCs, depending on the number of synchronous SNR inputs. The time course of firing rate increase was different in models fit to parkinsonian conditions due to reduced M-type and increased BK potassium conductances (Fig. 5). Beta frequency (12.5 or 25 Hz) modulations in SNR inputs were sufficient to induce spike-phase locking in VM (Fig. 6). The resulting beta modulation depth of VM spiking was amplified by the VM transfer function and was stronger than that of the SNR inputs. Adding beta oscillations in DRIs enhanced this effect (Fig. 7). Therefore, VM TCs relayed and amplified the oscillatory patterns in DRIs and SNR inputs (Fig. 6, 7), without differences between normal and parkinsonian states.

### Effects of dopamine depletion in vitro

In vitro recordings showed that dopamine depletion increased excitability of VM as a result of M-type current suppression (Bichler *et al*., 2021). Comparing the parameters of normal and parkinsonian models, we determined differences in their T- and M-type currents. In particular, the M-type conductance was 26-fold lower in parkinsonian models than control, and a subsequent 32-fold increase in M-type conductance yielded f-I curves and rebound spike counts in parkinsonian models indistinguishable from control. M-type current in thalamus is normally regulated by acetylcholine, where the activation of muscarinic receptors depolarizes thalamic neurons during high-attentional states (Llinás and Steriade, 2006). The main source of VM cholinergic input is from the pedunculopontine tegmentum and the laterodorsal tegmental nucleus (Huerta-Ocampo *et al*., 2020), and optogenetic stimulation of cholinergic neurons in this area switches sleep states from NREM to REM in mice (Van Dort *et al*, 2014). Indeed, optogenetic activation of VM neurons directly promotes arousal and wakes mice from NREM sleep (Honjoh *et al*., 2018). Therefore, it is plausible that M-type current downregulation in VM thalamus would be causative of disturbed sleep states in Parkinson’s disease that are characterized by insomnia, increased arousal and restless leg syndrome (for review, see (Lajoie *et al*., 2021)). We speculate that, in agreement with our simulations, muscarinic positive allosteric modulators that enhance M-type (Kv7) currents may have anti-parkinsonian effects on motor thalamic activity in PD and could improve associated sleep disorders.

### Effects of dopamine depletion on thalamic firing rates in vivo

Our results indicated that firing rates in response to simulated in vivo input conditions in the parkinsonian VM models were robustly increased compared to control models because of M-type potassium current reduction. However, robust rate increases are not observed in thalamic recordings from dopamine-depleted rodents in vivo (Anderson *et al*., 2015; Di Giovanni *et al*., 2020; Nakamura *et al*., 2021). As classic models suggested that motor deficits in PD might be due to increased basal ganglia inhibition to motor thalamus, and that this might cause motor deficits (DeLong, 1990), we tested the hypothesis that increased inhibition in conjunction with increased excitability could lead to a normalization in firing rates. Indeed, increasing the conductance of SNR synapses by 15% yielded firing rates in parkinsonian models indistinguishable from control. Therefore, increased excitability of VM neurons after dopamine depletion might be a compensatory mechanism for increased inhibition in this state.

Our sensitivity analysis of the effect of different input sources on VM firing suggested that VM activity in vivo was primarily influenced by SNR inhibition, whereas the impact of RTN inhibition was weak (Fig. 4). This was primarily due to the smaller size and more distal location of RTN terminals that we adopted from the findings of anatomical studies (Ilinsky and Kultas-Ilinsky, 2002; Rovó *et al*., 2012). This is consistent with experimental findings showing that the impact of non-nigral inhibition on VM of mouse was not substantial (Chevalier and Deniau, 1982).

### Impact of nigral bursting on motor thalamic activity

In vitro studies demonstrated that synchronous SNR activation could evoke rebound bursting based on T-type Ca^2+^ current activation in VM (Edgerton and Jaeger, 2014b). However, these experiments were carried out in the absence of background in vivo excitatory inputs that will limit the hyperpolarization achieved by nigral bursts. This effect of excitatory inputs was borne out by our simulations, where even 100% of synchronous SNR bursting in the presence of a tonic background of inhibition and excitation did not allow the development of T-type rebound bursts (Fig. 5). Instead, we found that under these conditions M-type and BK potassium currents caused a slower post-inhibitory firing increase in VM neurons that was correlated with the percentage of synchronous SNR inputs (Fig. 5). Moreover, in our simulations, identical SNR inputs evoked rebound spiking with different courses in normal and parkinsonian states (Fig. 5), due to changes in M-type and BK current activation. This suggests that an altered course of post-inhibitory spiking could be a marker of parkinsonian pathophysiology in VM, consequent to the homeostatic changes induced by dopamine depletion.

### Impact of beta oscillations in the afferent inputs to VM

In vivo recordings showed that dopamine depletion enhanced beta oscillations in motor areas and that VM promoted their propagation to motor cortex (Brazhnik et al., 2012; Brazhnik et al., 2016). We simulated beta oscillations at 12.5 Hz or 25 Hz as a spike-phase locking in the activity of SNR input to VM, and/or cortical MODs and DRIs input, consistent with the experimental recordings (Brazhnik et al., 2012; Brazhnik et al., 2016). Particularly, recordings showed that beta band oscillations were prevalent in non-attentive resting states whereas increased beta oscillations occurred during treadmill walking (Brazhnik *et al*., 2012). In our simulations, low-amplitude beta oscillations in the afferent inputs resulted in significant spike-phase locking in VM at both 12.5 and 25 Hz (Fig. 6, 7), which not only relayed but also amplified the oscillatory patterns in input (Fig. 6, 7). This amplification was not different between normal and parkinsonian conditions. Therefore, our simulation results suggest that the increased beta oscillations and spike-phase locking observed in dopamine-depleted animals (Brazhnik *et al*., 2016; Nakamura *et al*., 2021) is a consequence of changes in the activity of the afferent inputs to VM and do not rely on intrinsic excitability changes in VM.

In vivo experiments showed the necessity of SNR-VM interactions for propagation of beta oscillations to motor cortex (Brazhnik *et al*., 2016). Our simulations showed that beta oscillations in SNR alone were sufficient to induce significant spike-phase locking in VM (Fig. 6), supporting the possibility that beta oscillations could originate in SNR and then entrain the cortico-thalamocortical loop through the VM. However, results in dopamine-depleted mice indicate a lack of beta frequency increases in SNR (Lobb and Jaeger, 2015; Willard *et al*., 2019), suggesting a species-specific or state-dependent outcome in this regard.

In conclusion, modeling motor thalamic synaptic integration in normal and parkinsonian states indicates that parkinsonian firing pattern changes in the inputs are readily transmitted to the outputs. Increased intrinsic excitability may primarily act to compensate for changes in the excitatory / inhibitory input balance and lead to a homeostatic recalibration of firing rates. The increased excitability may come at the cost of accurate cholinergic modulation in motor thalamus, however, and contribute fluctuations in cortical arousal.

## Acknowledgments

The authors thank Taylor M. Kahl for helping in proofreading the manuscript. Simulations were performed using the Neuroscience Gateway (NSG) resource and an XSEDE cluster allocation (NAT220001) granted to FC. This work was supported by NINDS grants P50-NS098685 and 1P50NS123103 to DJ.

